# Fitness Landscapes Reveal Simple Strategies for Steering Evolution to Minimize Antibiotic Resistance

**DOI:** 10.1101/093153

**Authors:** Maria Smith, Sarah Cobey

## Abstract

The evolution of antibiotic resistance presents a practical and theoretical challenge: the design of strategies that limit the risk of evolved resistance while effectively treating current patients. Sequentially cycling antibiotics has been proposed as a way to slow the evolution of resistance by reducing the extent of adaptation to a given drug, and clinical trials have demonstrated its effectiveness in some settings. Empirical fitness landscapes in theory allow the sequence of drugs to be refined to maximize tradeoffs between drugs and thereby slow adaptation even further. Using the measured growth rates of 16 genotypes of *Escherichia coli* in the presence of *β*-lactam antibiotics, we test an adaptive strategy, based on a Markov chain transition matrix, to select drug sequences that continuously minimize resistance. Cycling is never selected over the long term. Instead, monotherapy with the antibiotic that permits the least growth in its landscape’s absorbing state is rapidly selected from different starting conditions. Analysis of a synthetic fitness landscape shows that cycling drugs that induce sensitivity to one other could, in theory, outperform monotherapy. These results underscore the importance of considering the specific topologies of fitness landscape in determining whether to cycle drugs and suggest a general computational approach to identify high performing, practical strategies to manage resistance.

## Introduction

Individual treatment decisions collectively impose strong selection for antibiotic resistance in many microbial populations. Various strategies have been suggested to slow this evolution. For instance, many patients receive high-dose antibiotic prescriptions that they are advised to complete regardless of symptoms. But the intended impacts of such policies can be far from theoretical expectations [1,2] and observed outcomes. Simple models suggest many conditions under which aggressive antibiotic treatment accelerates the emergence of resistant populations, for instance, if antibiotics release resistant strains from competition with sensitive strains [2] or create temporal or spatial gradients in drug concentration to ease adaptation [3,4]. Effective strategies to manage resistance thus hinge on accurate assumptions about microbes’ ecological and evolutionary dynamics [5].

One proposed method to limit the evolution of resistance, especially in healthcare settings, is antibiotic cycling [6–12]. The idea is to treat all patients in a hospital ward with the same antibiotic for a given period, and then with a second antibiotic, and so on, eventually cycling back to the first. Sequential monotherapy might reduce the probability that microbes become resistant to more than one antibiotic at a time, especially if drugs are switched before or soon after resistance is detected. When multidrug resistance is not already common and microbial populations are well mixed [13,14], this approach is expected to work [8,15], and a recent meta-analysis showed that cycling has been effective in reducing resistance and disease burden in several pathogens in hospitals [8].

Cycling could be even more effective if the sequence of drugs were chosen to minimize the probability of accumulating resistance to sequential drugs [9–11,16,17]. In evolutionary terms, these sequences would exploit known tradeoffs: given emergent resistance to drug A, choose drug B so that resistance to drug B is difficult (or impossible) to attain. Tradeoffs may be weak to nonexistent. For instance, resistance to drug A might involve the acquisition of an efflux pump, which could be effective against drug B. In this case, known as cross-resistance [18–20], there is no advantage to cycling. In a phenomenon known as negative cross-resistance or collateral sensitivity, resistance to drug A requires sensitivity to drug B [21,22]. One of the best examples of such adaptive tradeoffs is the evolution of resistance to a class of antibiotics, the *β*-lactams, in *Escherichia coli* [23–29]. Resistance to *β*-lactams is chiefly determined by the *β*-lactamase gene, and different alleles are associated with different degrees of resistance to specific drugs. No single allele confers maximal resistance to all *β*-lactams. Thus, drugs might be ordered in such a way to increase the number of mutations required to gain resistance to one drug, assuming adaptation to the previous one. If required intermediate mutations decrease fitness, they might not be selected, effectively blocking the evolution of resistance to the next drug. Adaptation can thus be prevented or greatly slowed [12,16,17,21].

In pursuit of this goal, the fitnesses of different *E. coli* genotypes were previously enumerated for each of 15 *β*-lactam antibiotics and used to identify optimal sequences for antibiotic cycling [9,25]. These fitnesses, measured by growth rates in the presence of each antibiotic, were obtained for the wild-type, “sensitive” genotype (known as TEM-1) and a more broadly resistant genotype (known as TEM-50) and all permutations (TEM-1 or TEM-50) at each of the four loci that differ between the genotypes, i.e., for 16 genotypes in total. These data describe a small subset [23, 30] of an evolutionary fitness landscape: genotypes differing by one amino acid mutation form a connected network in genotype space, and each genotype has a fitness (growth rate) in each environment (antibiotic). Fitness landscapes allow potential evolutionary trajectories to be modeled explicitly[31]. From any starting genotype, the probability of transitioning to a neighboring genotype can be calculated from neighbors’ relative growth rates. When antibiotics are switched, more fit genotypes might suddenly become less fit, decreasing resistance.

Enumeration of fitness landscapes is a well-known challenge to the development of effective strategies to manage antibiotic resistance [22,24,26,27,29,32–36]. It is undoubtedly useful to know which sequences of antibiotics facilitate or hinder the acquisition of resistance, especially if current practice unknowingly promotes the former. We show here that the way in which these landscapes are analyzed is equally important. Using the *β*-lactam landscapes of *E. coli* as a proof of concept, we first compare current approaches to select optimal antibiotic sequences for cycling. We show that the extent of adaptation allowed under each antibiotic and the number of times a cycle is repeated have large impacts on which antibiotic sequence is recommended and on the efficacy of antibiotic cycling in general. We next consider a practical challenge of cycling: under some criteria, “optimal” antibiotic sequences can involve periods of high resistance. We investigate what would happen if antibiotics were chosen adaptively, with the aim of using the antibiotic that minimizes bacterial growth given the current distribution of bacterial genotypes. One particular antibiotic consistently arises in these cases: cycling never helps. We show why, partly by demonstrating with a synthetic fitness landscape how cycling sequential drugs could maximize collateral sensitivity. Taken together, our results demonstrate that modeling evolution on fitness landscapes can inform policies to minimize resistance.

## Methods

### Empirical fitness landscapes

Empirical fitness landscapes were used to model the possible paths of evolution in multiple environments, each defined by the administration of a single antibiotic. The landscape is a topology of all possible permutations of mutations in a set of genetic loci, and genotypes are connected if they are a single amino acid mutation apart.

In *E. coli,* the wild-type *bla*_TEM−1_ gene coding for *β*-lactamase needs four amino acid substitutions to become *bla*_TEM−50_, which is resistant to cephalosporins and inhibitor combined therapies. Mira et al. [9] created all 16 possible genotypes and measured their growth rates as proxies of fitness under 15 different *β*-lactam antibiotics, creating 15 different fitness landscapes. Each genotype has a range of growth rates depending on the antibiotic, suggesting the potential for successful reversion of resistance given the right antibiotic sequence (Table S1).

### Markov model for evolution

For each antibiotic treatment, the growth rates were organized into directed fitness graphs, where genotypes *u* and *v* are directionally connected (*u → v*) if and only if they differ by a single mutation and *v* has a higher fitness than *u* (Fig. 1) [9]. Each fitness graph under an antibiotic, *A*, was mapped to a 16 × 16 transition matrix *M*(*A*) with rows and columns corresponding to all possible genotypes. For genotypes *u* and *v*, the entry at row *u*, column *v* of *M(A)* is the probability of transitioning from genotype *u* to *v* after treatment with antibiotic *A*. It is given by

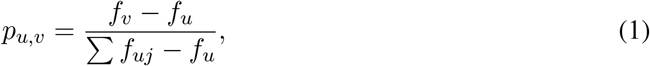

where *f_v_* and *f_u_* are the growth rates of genotypes *v* and *u* and *uj* are adjacent genotypes to *u* with higher growth rates. If genotype *u* is not adjacent to *v* in the fitness graph or *v* has lower fitness than *u*, *p_u,v_* = 0. Equation 1 indicates that as *f_v_* increases with respect to the fitness of other adjacent genotypes of *u*, *p_u,v_* increases, so transitions to higher fitness genotypes are more likely. A genotype *u* with no fitter neighbors has *p_u,u_* = 1.

**Figure 1:**
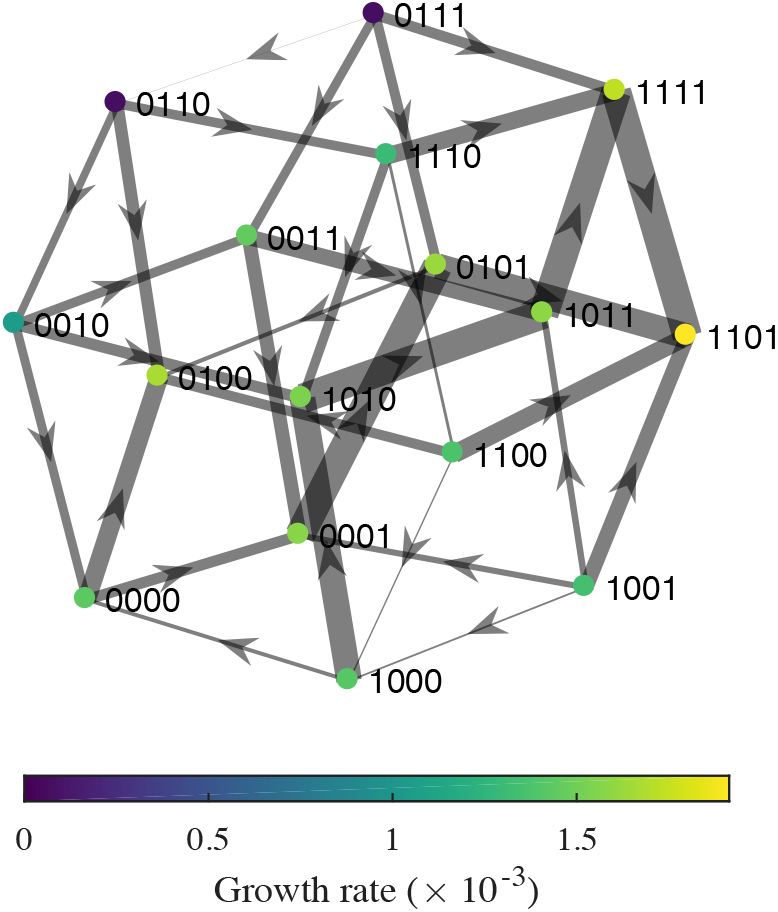
Fitness graph for amoxicillin-clavulanic acid (AMC). Node colors show genotype growth rates. Edge weights show the relative transition probabilities *p_u,v_* from each source node *u*. Edges are not shown for transitions with zero probability. Self-loops (*p_u,u_* = 1) are omitted for clarity.

We use these transition probabilities to approximate evolution under successive antibiotic treatments. Let *M* (*A_i_*) be the transition matrix for antibiotic *i* given by the fitness graph *A_i_*. If *A*_1_, *A*_2_,…, *A_n_* is a sequence of antibiotics, the entry in row *u*, column *v* of the matrix obtained by matrix multiplication of the transition matrices

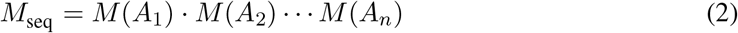

gives the net probability of transitioning from genotype *u* to genotype *v* at the end of the antibiotic cycle.

Assuming an initially sensitive genotype 0000, the vector of initial genotype densities is given by

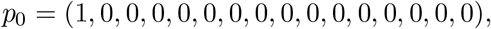

and the equation

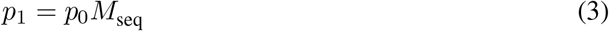

gives the frequency distribution *p_1_* for being in each genotype after one application of the antibiotic sequence *A*_1_, *A*_2_,… , *A_n_*. Implicitly, this approach assumes the microbial population has little time to adapt to each drug before it is switched: genotypes mutate to all possible first-degree neighbors but no further, and these new mutants coexist in proportion to their relative fitnesses. We refer to antibiotic treatment in this scenario as having a “short period”.

### Effects of long-term cycling, treatment length, and immigration

One optimization strategy is to calculate the probability of returning to the initial sensitive genotype after every possible sequence of *k* unique antibiotics and then select the sequence that maximizes the frequency of the sensitive genotype [9]. If *M*_seq_ is the transition matrix for a single cycle of an antibiotic sequence, then (*M*_seq_)*^n^* is the matrix giving the transition probabilities after *n* cycles, and

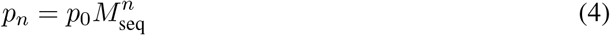

gives the genotype frequency distribution *p_n_* after *n* cycles of the sequence. Taking *n → ∞*, we can find the long-term equilibrium frequencies of each genotype, including the sensitive genotype.

Another approach is to assume that pathogen strains fully adapt before antibiotics are switched. Nichol et al. [10] assume that the microbial population reaches evolutionary equilibrium under each antibiotic and finds *M*_seq_by multiplying the equilibrium distributions for each individual antibiotic in the sequence:

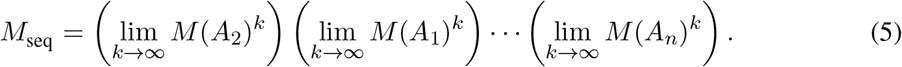

Because treatment with each antibiotic implicitly lasts infinitely long (i.e., allows maximal adaptation), we call this the “long period” scenario. Note that since matrix multiplication is not commutative, *M*_seq_ given by equation 5 is not equivalent to the 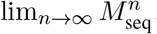, where 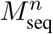 is the transition matrix from equation 4. In other words, the set of long-term, equilibrium transition probabilities after infinite cycles does not equal the set of transition probabilities obtained from a single cycle, allowing maximum adaptation to each antibiotic.

These approaches so far assume that mutations occur incrementally, or that the only accessible genotypes are one amino acid mutation away. The approaches also only allow transitions that increase fitness. We can instead optimize drug selection while assuming that multiple simultaneous mutations (or immigration of genetically distant strains) and transitions to less fit genotypes sometimes occur. In these scenarios, rather than assuming that the probability of transitioning to a less fit or non-adjacent genotype is 0, we allow any genotype to arise. Specifically, we let *q_u,v_* be the probability of transitioning from *u* to *v* with an immigration probability *p*_imm_ given by

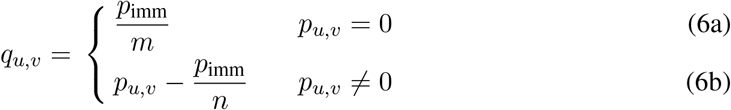

where *m* is the number of neighbors *v* such that *p_u,v_* = 0 (normally inaccessible genotypes) and *n* is the number of *v* such that *p_u,v_* ≠ 0 (normally accessible genotypes).

### An adaptive optimization strategy

Previous models have selected optimal sequences of drugs to cycle by maximizing the probability of returning to the sensitive genotype after a sequence of *k* antibiotics [9,10]. In healthcare settings, it is impractical to optimize long-term outcomes (the probability of returning to a sensitive genotype at the end of the cycle) if the short-term effects include high infection rates or resistance. If the aim is to minimize the incidence of bacterial infection constantly over time, a logical approach is to identify at every time step the antibiotic that minimizes the expected growth rate of the bacterial population. This is similar to the proposed “informed switching” policy [37] or the adjustable cycling approaches [8] proposed for individuals. We apply this motivation to steer evolution on fitness landscapes. We assume that switching to a new antibiotic occurs whenever the mean growth rate exceeds a threshold and can be lowered, given the current distribution of genotypes. Some antibiotics may thus have short periods and others long periods under this strategy, depending on the number of times they are applied before switching. There is no guarantee that cycles will arise.

## Results

### Long-term cycling and treatment length, but not immigration, affect strategy

We first investigated whether existing cycling strategies converge on similar results, especially when their assumptions are relaxed. Previously, antibiotic sequences were identified that give the highest probabilities of selecting sensitive strains after a single cycle [9] (Table S2). We considered the frequency of the sensitive genotype after infinite cycles, the theoretical limit if cycling were common. Instead of optimizing drug sequences for just one cycle, we optimized for an infinite number of cycles using the long-term equilibrium transition matrix (equation 4). The frequency of the sensitivity genotype tends to decline rapidly as cycles repeat (Fig. S1). Notably, rather than a 70% frequency of the sensitive genotype after one cycle with two drugs (*k* = 2), the best two-antibiotic sequence after infinite cycles yields only an 8% frequency of this genotype and involves different antibiotics (Table S2). With infinite cycles, longer sequences involving more unique antibiotics are required to maintain the frequency of the sensitive genotype, which peaks at 60% at the longest sequence tested (*k* = 5; Table S2). To determine if generally the same drug sequences would be selected independent of assumptions, for each scenario we ranked sequences of all lengths by the final frequency of the sensitive genotype. There is a negative correlation between sequences’ rankings after one cycle versus infinite cycles (Spearman’s *ρ* = −0.45, *p* < 0.001; Fig. 2a), demonstrating that the duration of cycling affects which sequence is recommended. Overall, compared to a single sequence of treatments, minimizing resistance with repeated cycles appears to require more drugs and to yield poorer outcomes (Table S2).

**Figure 2:**
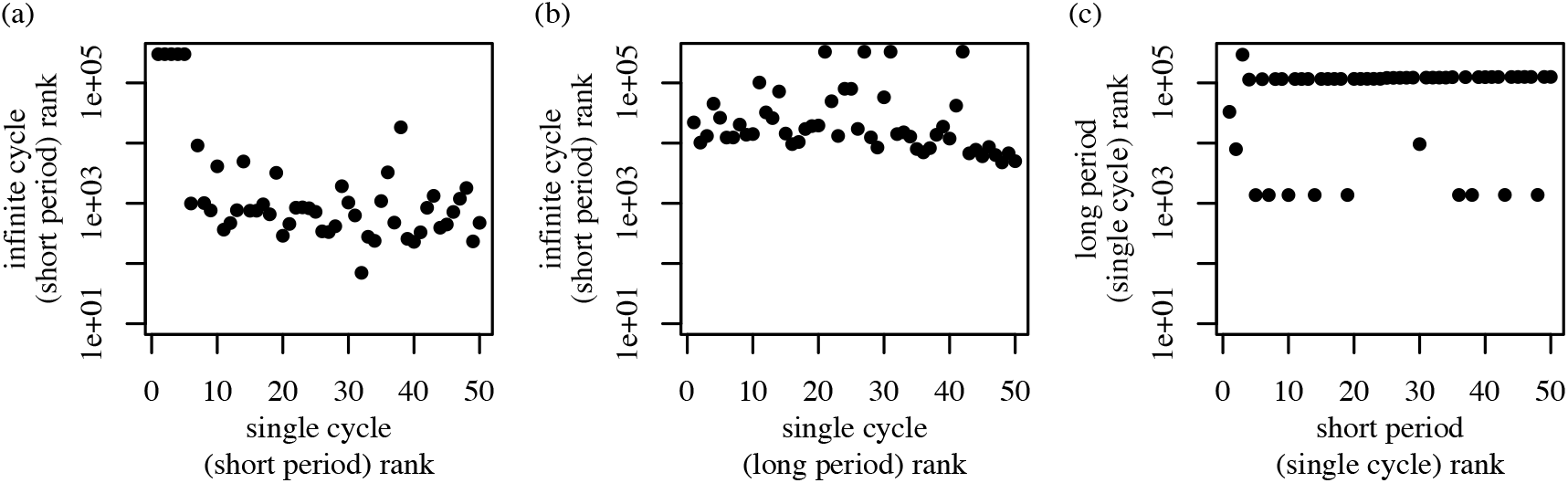
Correlations between cycle rankings in different scenarios. All sequences of lengths *k* = 2,3,4, 5 were ordered by the final frequency of the sensitive genotype, from highest to smallest. Ranks of the top 50 sequences in one scenario were compared to the same sequences’ ranks in another scenario. (a) Rankings with single [9] and infinite cycles. (b) Rankings with infinite cycles and short durations compared to single cycles and infinite treatment duration [10]. (c) Single-cycle rankings for short and infinite treatment durations.

The effective duration of treatment might also influence the optimal strategy because longer treatment periods allow more adaptation to each antibiotic. Several studies have suggested that cycling becomes less effective with longer periods, as a long cycling period effectively becomes single-drug treatment [8, 13]. In our model, long periods allow genotypes to converge to a stationary distribution under each drug before switching (equation 5). Considering the sequence of amoxicillin and pipercillin + tazobactam as an example, applying long periods leads to faster evolution away from the sensitive type than single applications of each drug (“short periods”) (Fig. S2). As expected, long periods select for different sequences of drugs. For tested sequence lengths *k* > 1, in no case is the optimal sequence the same for short and long periods (Table S2). A sequence’s rank with long periods and a single cycle and its rank with short periods and infinite cycles are negatively correlated (Spearman’s *ρ* = −0.43, *p <* 0.002; Fig. 2b). Considering single cycles alone, however, there is a positive correlation between sequences chosen for short and long periods (Spearman’s *ρ* = 0.60, *p <* 10^−5^; Fig. 2c). Thus, the optimal antibiotic sequence is sensitive to the extent of adaptation allowed under each drug and the number of cycles.

All of these approaches assume that mutations occur incrementally, or equivalently, that genotypes can only transition to adjacent states with higher fitness. Genotypes that are distant or less fit might arise from mutation or immigration and could also influence evolution. Allowing small (1%) total probabilities of transitioning to less fit and nonadjacent states often leads to different optimal strategies (Tables S2 and S3). Overall, however, these assumptions do not affect the predicted genotype frequency nearly as much as the choice of cycle number and period length (Fig. 3). Starting from the sensitive genotype, sensitivity is easiest to maintain when periods are short and cycles do not repeat. These results suggest a need to identify the optimal duration for each treatment and strategies that are more robust over time.

**Figure 3:**
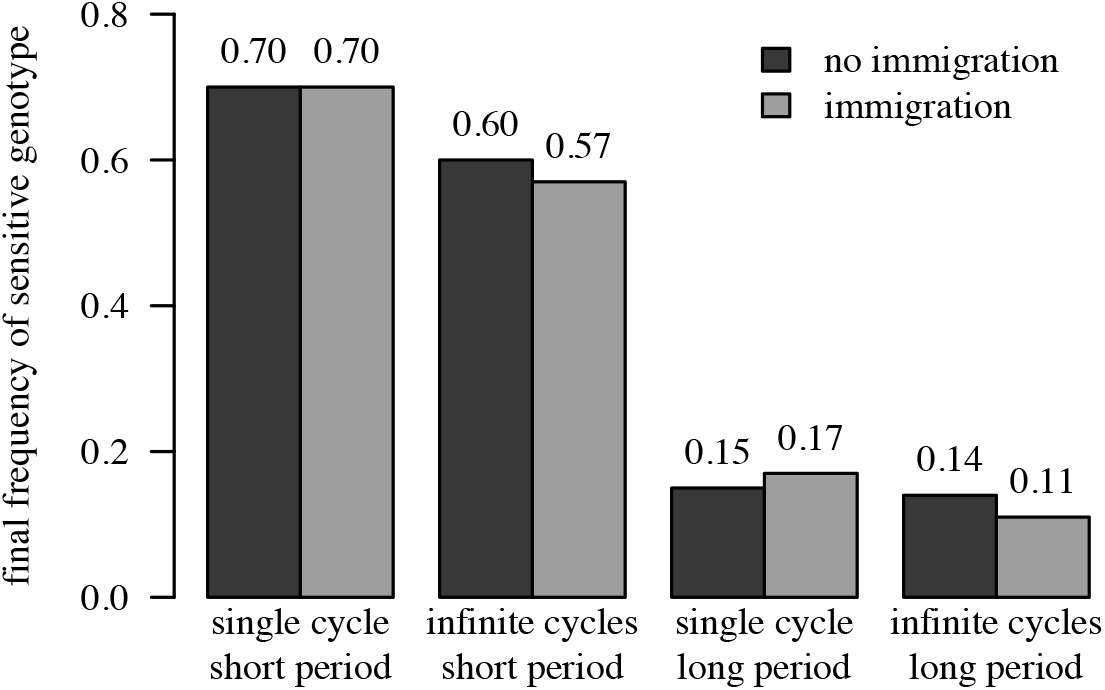
The final fraction of the sensitive genotype under different optimization strategies. Sequences of lengths *k* = 2,3,4, 5 were pooled for each of the four strategies with and without immigration.

### An adaptive strategy to minimize resistance

Strategies that minimize resistance in the long run may be impractical if they involve short-term costs: hospitals cannot sacrifice patients in May to improve outcomes in December (e.g., [38]). A more defensible approach is to select the antibiotic that minimizes the expected growth rate given the current population. Since finely monitoring genotype distributions is infeasible, we assume that the drug is reevaluated only when the mean growth rate exceeds a threshold, for which the frequency of failed treatments might be a proxy. The so-called sensitive genotype in fact has high growth rates with some drugs (Table S1), and thus optimal strategies might favor different genotypes. Under an adaptive approach, the optimal treatment duration of each antibiotic and any cycles arise endogenously.

We chose drugs adaptively, selecting the antibiotic that minimized the expected growth rate at each step, or continuing the current antibiotic if the growth rate remained below a threshold (here, 0.001). To facilitate comparison with the previous approach, we first assumed the sensitive genotype starts at 100% frequency. The adaptive strategy immediately selects for a dual-regimen monotherapy, amoxicillin + clavulanic acid (AMC) (Fig. 4a). This drug is also selected as long-term monotherapy when starting from other genotypes, although other drugs are sometimes chosen in an initial transient phase (Fig. 4b,c). Random initial conditions also eventually select for AMC (Table S4). Notably, the equilibrium genotype frequencies and thus equilibrium growth rates under long-term AMC therapy vary with the starting condition, ranging from 1.67 × 10^−3^ to 1.88 × 10^−3^. The former case arises when the population starts at genotype 0100, a local fitness peak under AMC (Fig. 1), and the latter when the equilibrium population is dominated by the other fitness peak, genotype 1101 (Table S5). Genotype 1101 has a higher growth rate than 0100 (1.91 × 10^−3^ v. 1.67 × 10^−3^) and a larger “basin of attraction,” in that it serves as the absorbing state for more genotypes (Fig. S3). Mean growth rates under the adaptive strategy are approximately 15% to 30% lower than the lowest growth rates under traditional antibiotic cycling (Table S2).

**Figure 4:**
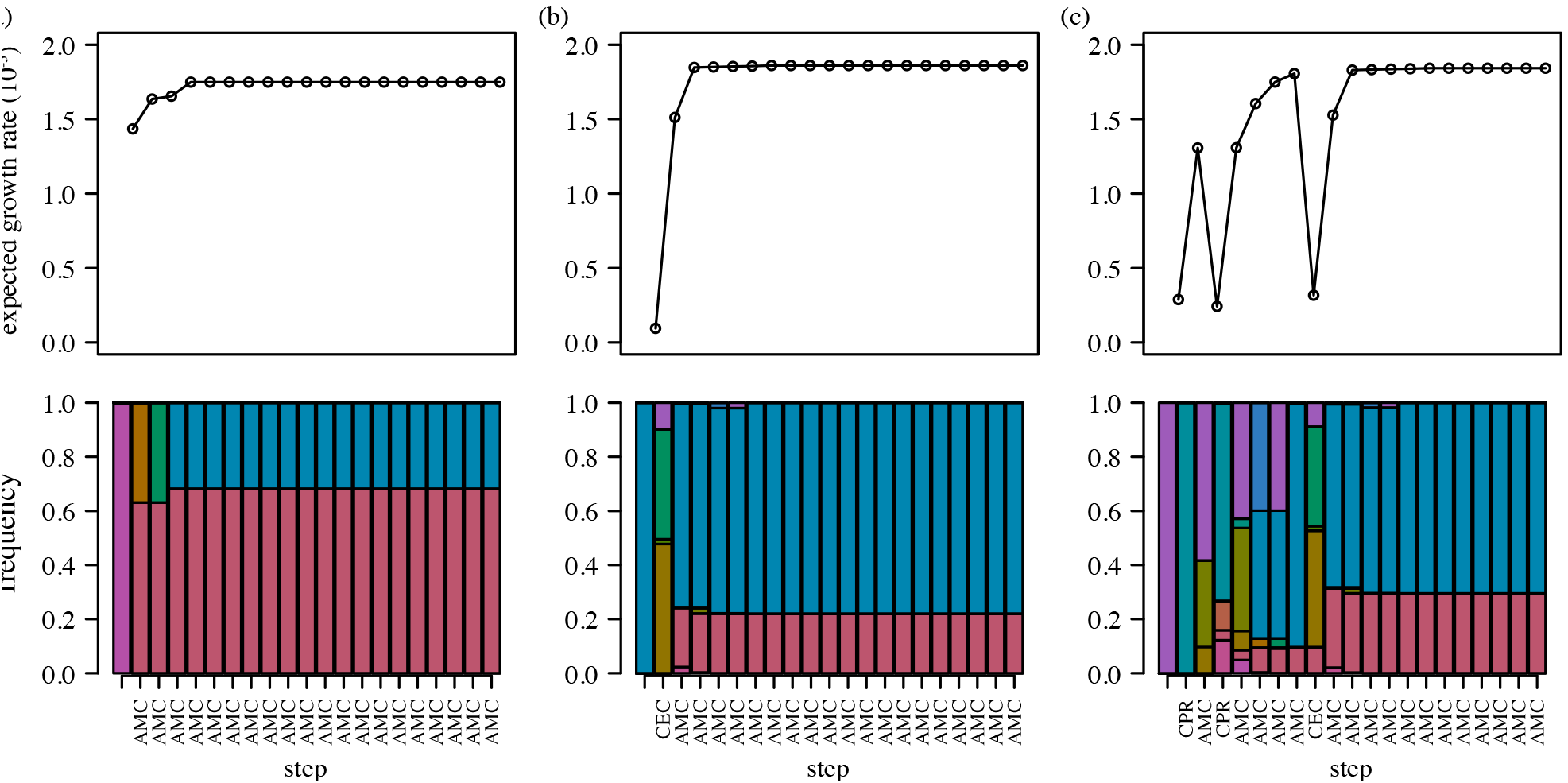
Sample time series starting from the (a) 0000, (b) 1101, and (c) 1111 genotypes under the adaptive strategy. The top row shows population growth rates corresponding to the genotype distributions below, with the selected drug shown for each step.

We investigated whether AMC monotherapy would suffice as a fixed regimen to apply regardless of starting conditions. Under the adaptive strategy, AMC monotherapy arises only after treatment with at least one other drug when starting from 10 of 16 individual genotypes. Applying AMC immediately in these cases leads to two types of cost. First, there is the potential cost of higher growth in the transient phase before equilibrium (e.g., during the non-AMC periods depicted in Fig. 4b,c and until the long-term growth rates have been attained). We defined the cumulative initial cost of the fixed AMC strategy as the cumulative difference in growth rates between the fixed and adaptive strategies until both are at equilibrium. Second, there may be persistent differences in growth rates due to shifts in genotypes’ limiting distributions as a result of the fixed therapy. We define the long-term cost as the expected difference in growth rates between the fixed and adaptive strategies at equilibrium.

As expected, both types of cost are present (Table S4). The cumulative transient cost is up to 128% of the mean equilibrium growth rate for single genotype initial conditions. From random starting conditions, the transient cost is low (0.12% of the mean equilibrium growth rate). Notably, fixed AMC therapy from single-genotype populations often leads to higher long-term frequencies of the most resistant genotype, 1101, than would appear had drugs been adaptively chosen (Tables S5 and S6). The higher frequencies increase the long-term growth rate as much as 3.89% over the adaptive strategy, although the average difference in growth rates between the two strategies for random initial populations is only 0.02%. Thus, when starting from heterogeneous populations, there is little cost to immediate AMC therapy. These results show that the effectiveness of a non-adaptive approach is sensitive to the initial genetic diversity.

### Cycling could be selected, in theory

Cycling should be advantageous if the fitness landscapes of different drugs are negatively correlated: relatively high-fitness genotypes in one environment have relatively low fitness in another. In other words, for neighboring genotypes *u* and *v*, we expect *u → v* under drug A and *u ←v* under drug B (equivalently, 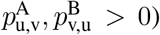. The two genotypes comprising the absorbing states under AMC, 0100 and 1101, in fact have lower fitness than their respective neighbors 0110 and 0101 under cefprozil (CPR), and the population thus transitions toward these genotypes when CPR is applied (Fig. S4). However, the growth rates of 0110 and 0101 under CPR are high enough that cycling leads to higher mean population growth rates (during the CPR phase) than if AMC alone were applied. This example reiterates that selection for antibiotic cycling requires not only that local fitnesses be negatively correlated but also that neighboring genotypes’ fitnesses be lower under the new drug. More precisely, the mean population growth rate must be less under the new drug than the previous drug after taking Markov transition densities into account.

To test this principle, we constructed a synthetic fitness landscape by reversing the direction of flow along the AMC fitness landscape (Figs. 1 and S3). Specifically, we ranked genotypes by their growth rates under AMC and then swapped the highest and lowest growth rates, the next highest and next lowest, and so on. We then added the synthetic drug, rAMC, to the original list of 15 drugs and reran the adaptive selection routine. Long-term adaptive cycling of AMC and rAMC always arises and leads to dramatically lower growth than monotherapy: mean growth rates cycle between 1.61 × 10^−3^ to 1.66 × 10^−3^, a 12% improvement (Fig. 5). Although the two landscapes are perfectly negatively correlated and have equally high peaks, these peaks are surrounded by lower-fitness genotypes that are repeatedly selected during cycling.

**Figure 5:**
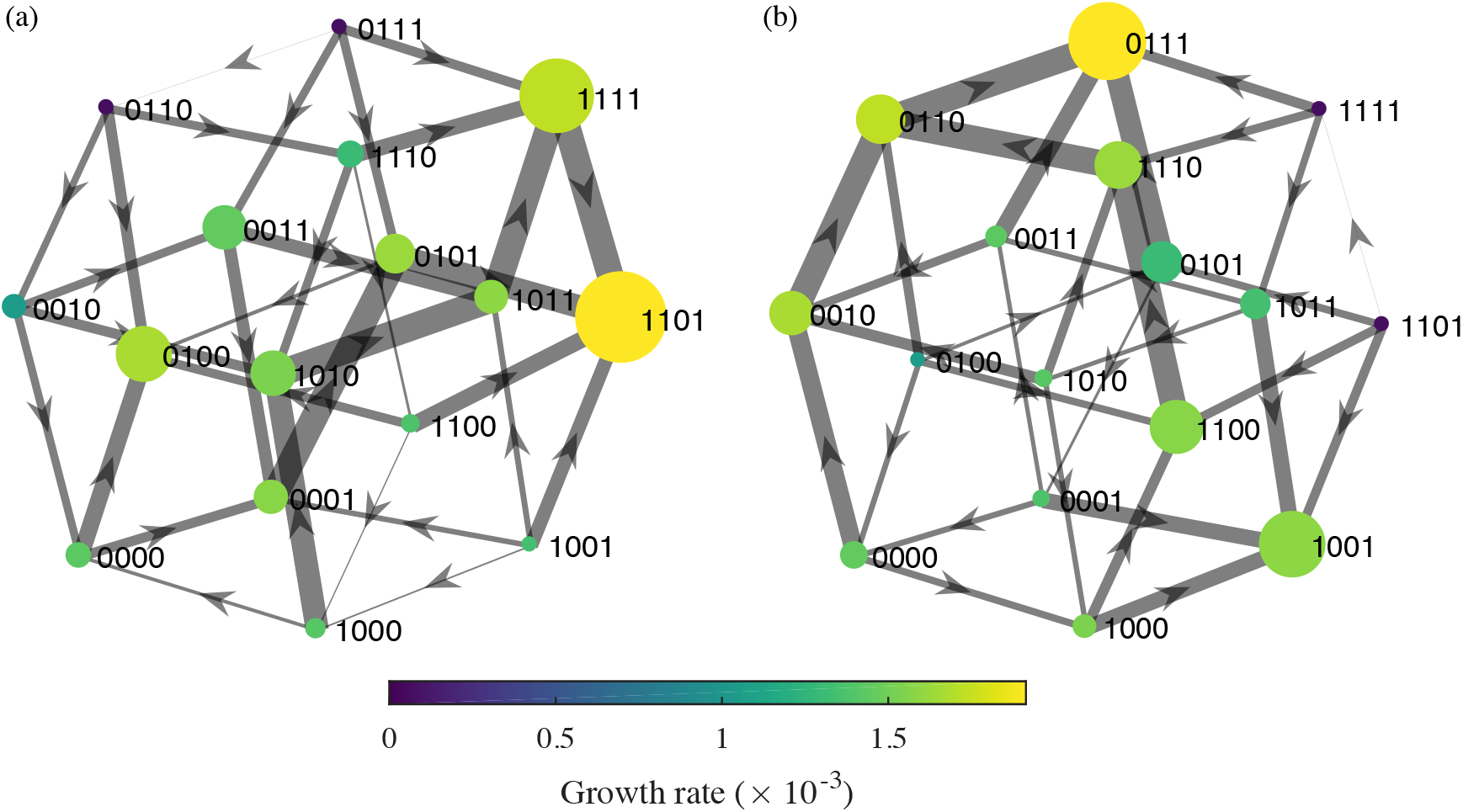
Fitness graphs during alternating (a) AMC and (b) rAMC treatments for the two-drug cycle at equilibrium. Node colors show genotype growth rates in each drug, node sizes indicate relative frequencies, and edge weights show the relative transition probabilities from each source node. Genotypes with near-zero frequencies are given visible mass to reveal color.

## Discussion

Antibiotic cycling belongs to a group of theoretical approaches that attempt to exploit the topologies of fitness landscapes to constrain the evolution of resistance [9,10,24,32]. We tested a simple, adaptive strategy that uses a Markov chain transition matrix to select drug sequences that minimize the expected growth rate of the pathogen population. When applied to the empirical fitness landscapes of *E. coli,* the adaptive strategy rapidly converges to therapy with a single drug. This result does not invalidate cycling as a potentially optimal strategy, which we demonstrate with a synthetic landscape, but it underscores the importance of quantitative assessments of empirical landscapes. We also posit that previous approaches to selecting antibiotic sequences for cycling are impractical in healthcare settings. Such strategies implicitly treat patients unequally over the course of the cycle, and they can thus lead to more resistance over time. Overall, our results suggest a simple approach for determining optimal treatments, which might include (but is not limited to) cycling antibiotics. We find that the long-term drug sequence that emerges under the adaptive approach can be applied as a fixed strategy with modest cost, suggesting that general recommendations may be possible.

The effects of AMC-rAMC cycling are similar to those of combined drug regimens that induce collateral sensitivity or cross-resistance [12,34], but here the tradeoffs are realized over time [17,39], and the selected strategy should in theory remain effective indefinitely. Collateral sensitivity occurs when the evolution of resistance to drug A simultaneously selects for sensitivity to drug B. In experiments, cycling drugs that induce collateral sensitivity slowed the evolution of resistance in *E. coli* [17] and *Staphylococcus aureus* [16]. Genotypes 1101 and 0101 demonstrate collateral sensitivity with AMC and CPR (Fig. S4), and more dramatic instances of collateral sensitivity can be found for other drug pairs (e.g., ampicillin-cefpodoxime with genotypes 1001-1011 and amoxicillin-cefaclor with 0101-1101) and for other genotype pairs with AMC-CPR (e.g., 0110-1110) (Table S1). In contrast to AMC and rAMC, which promote pervasive collateral sensitivity, cycling with between these other drug pairs is not selected over the long term. Only some treatments featuring collateral sensitivity also accommodate the need to minimize short-term pathogen growth rates. This result underscores the utility of studying the topology of fitness landscapes, not simply the rates of resistance to different drugs or correlations in drug sensitivities, in determining effective treatments to minimize the evolution of resistance.

The robustness of the AMC strategy to different starting conditions is a property of the fitness landscape. Notably, the genotype frequencies do not converge to an identical stationary distribution regardless of starting conditions. However, the genotype frequencies reach a small set of limiting distributions because the Markov chain, defined by *M*(*A*_AMC_), contains two absorbing states (Fig. S3). These states correspond to relatively low fitness peaks at genotypes 0100 and 1101. Because the graph forms a single component (all nodes are connected by at least one edge), at least one absorbing state is reachable from any starting condition. If the graph contained multiple components, the final genotype distribution could be especially sensitive to initial conditions, and it is possible that different long-term drug sequences would be optimal for some of them. General strategies may not exist in these cases (or even in cases involving a single component), but they raise opportunities for highly effective control if one drug selects for a subset of genotypes that cannot adapt to the second drug. A practical consideration is whether the persistence of less fit genotypes or multiple simultaneous mutations in nature might effectively connect otherwise unconnected subgraphs, allowing resistance to evolve.

The performance of the adaptive approach might be improved by considering biological “details” that the model omits, and such investigations are important before extending the results to clinical practice. The Markov chain is a radically simplified approach to evolving competing populations on an already simplified adaptive landscape. A strong assumption is that a single treatment lasts as long as it takes genotypes to mutate to all neighbors (and no further), and that mutants’ abundances scale linearly with their normalized growth rates (equation 1). In practice, possible mutations might not be realized, realized mutations will not be synchronous, and competition often leads to nonlinear growth. The model effectively assumes weak competition for resistance in a given environment and instantaneous transitions (with strong selection) between environments; both assumptions have dynamical consequences [9,40,41]. The model also omits host population structure. Others have shown that resistance can evolve more slowly if antibiotic use is heterogeneous among hosts [13,14]. Departures from sequential monotherapy will be required if the selected antibiotics cannot be tolerated by everyone. If entire subpopulations (e.g., children, immunocompromised patients, etc.) are exempted from some drugs, and these populations make unique contributions to transmission, then multiple fitness landscapes and a dynamical transmission model may be required to predict evolution accurately [8,37]. Finally, the landscapes themselves might change from the appearance of compensatory mutations, linked resistance elements, and previously undetected epistasis and pleiotropy [23,28,34,40,42]. It has been suggested that the *β*-lactamase mutations observed in nature most resemble those under fluctuating selective pressures in vitro, and mutations that appear in monotherapy in vitro are rarely observed outside the lab [43]. The distribution of mutations in nature might thus make an unstable basis for inferring the true breadth of viable mutations, and new strategies could expose new parts of the fitness landscape.

The effectiveness of antibiotic cycling in managing resistance to different pathogens in different settings is still in early stages of investigation, but the approach shows promise [8,16,17]. We have proposed a new and general strategy to select antibiotic sequences that takes advantage of data on empirical fitness landscapes, and we identify a specific treatment that may be effective for managing *β*-lactam resistance in *E. coli.* Considering heterogeneity in the host population and complexity in mutational and competitive dynamics is an important prerequisite for applying these recommendations. Although a strength of the adaptive strategy is that it prioritizes resistance reduction equally across patients in time, it does not solve one conflict inherent to almost all antibiotic use [1,21]: the most effective treatment for an individual patient is not always the one that minimizes resistance in the population. For instance, the most inhibitory drug for a patient infected with the 1101 genotype is cefotaxime. If the original drug works well enough, however, then this conflict might be avoided.

## Acknowledgements

This work was completed in part with resources provided by the University of Chicago Research Computing Center. We thank Stefano Allesina, Lauren Childs, Jacopo Grilli, Marc Lipsitch, and Pleuni Pennings for comments.

## Author contributions

Conceived and designed the experiments: MS SC. Performed the experiments: MS SC. Analyzed the data: MS SC. Wrote the paper: SC.

## Software

The code used in this analysis is publicly available at github.com/smithmj/Antibiotic-Cycling.

## Supporting Information

**Figure S1:**
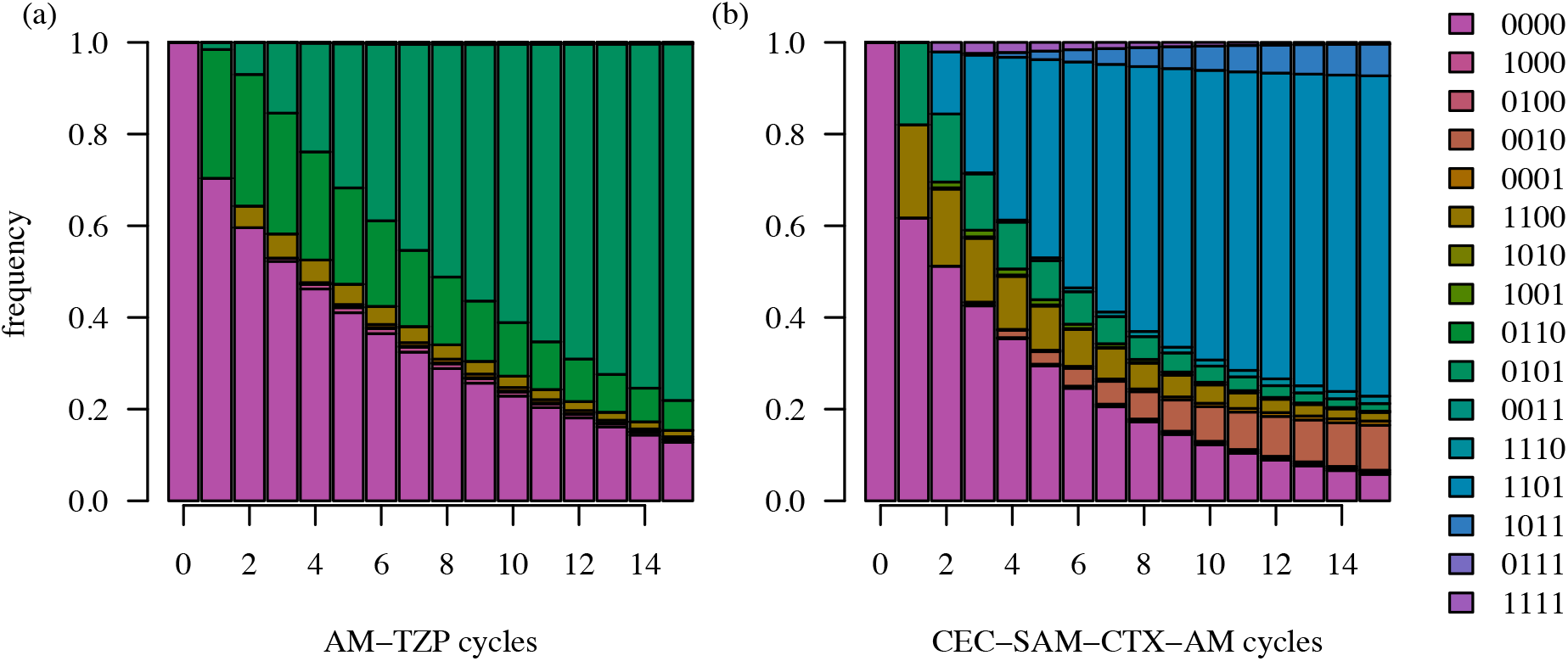
Evolution after multiple antibiotic cycles for two of the optimal sequences determined for single cycles with short periods [9].

**Table S1:** Growth rates (×10^−3^) of sixteen genotypes in the presence of fifteen drugs, reprinted from [9], The four-digit column labels give each genotype, with 0000 representing the sensitive wild-type *bla*_TEM-1_ and 1111 the more broadly resistant *bla*_TEM-50_ genotype. AMC is highlighted in gray.

|  | 0000 | 1000 | 0100 | 0010 | 0001 | 1100 | 1010 | 1001 | 0110 | 0101 | 0011 | 1110 | 1101 | 1011 | 0111 | 1111 | mean |
| --- | --- | --- | --- | --- | --- | --- | --- | --- | --- | --- | --- | --- | --- | --- | --- | --- | --- |
| Amoxicillin (AM) | 1.78 | 1.72 | 1.45 | 2.04 | 1.78 | 1.56 | 1.80 | 2.01 | 1.18 | 1.54 | 1.75 | 1.77 | 2.25 | 2.01 | 0.06 | 2.05 | 1.67 |
| Amoxicillin/clavulanic acid (AMC) | 1.44 | 1.42 | 1.67 | 1.06 | 1.57 | 1.38 | 1.54 | 1.35 | 0.07 | 1.63 | 1.46 | 1.31 | 1.91 | 1.59 | 0.07 | 1.73 | 1.32 |
| Ampicillin (AMP) | 1.85 | 1.57 | 2.02 | 1.95 | 2.08 | 2.19 | 0.05 | 2.17 | 2.03 | 2.20 | 2.43 | 0.09 | 2.32 | 0.08 | 0.03 | 2.82 | 1.62 |
| Ampicillin + sulbactam (SAM) | 1.88 | 2.20 | 2.46 | 0.13 | 2.53 | 2.50 | 2.31 | 2.57 | 0.08 | 2.43 | 0.09 | 2.53 | 3.00 | 2.89 | 0.09 | 3.45 | 1.95 |
| Cefaclor (CEC) | 2.26 | 0.23 | 2.40 | 2.15 | 2.00 | 2.15 | 2.24 | 0.17 | 2.23 | 1.85 | 2.65 | 2.64 | 0.19 | 0.09 | 0.21 | 0.52 | 1.49 |
| Cefepime (FEP) | 2.59 | 2.07 | 2.44 | 2.39 | 2.57 | 2.74 | 2.96 | 2.45 | 2.65 | 2.81 | 2.83 | 2.80 | 2.86 | 2.63 | 0.61 | 3.20 | 2.54 |
| Cefotaxime (CTX) | 0.16 | 0.19 | 1.65 | 1.94 | 0.09 | 0.23 | 1.97 | 0.14 | 2.30 | 0.14 | 2.35 | 0.12 | 0.09 | 0.20 | 2.27 | 2.41 | 1.01 |
| Cefotetan (CTT) | 2.13 | 3.24 | 3.29 | 2.80 | 1.92 | 0.55 | 2.88 | 2.97 | 3.08 | 2.89 | 0.59 | 3.19 | 3.18 | 0.89 | 3.51 | 2.54 | 2.48 |
| Cefpodoxime (CPD) | 0.60 | 0.43 | 1.76 | 2.60 | 0.25 | 0.64 | 2.65 | 0.39 | 2.91 | 1.47 | 3.04 | 0.96 | 0.99 | 1.10 | 3.10 | 3.27 | 1.63 |
| Cefprozil (CPR) | 1.74 | 1.55 | 2.02 | 1.76 | 1.66 | 0.22 | 0.17 | 0.26 | 2.04 | 2.05 | 1.79 | 1.81 | 0.24 | 0.22 | 0.22 | 0.29 | 1.13 |
| Ceftazidime (CAZ) | 2.13 | 0.29 | 2.04 | 2.62 | 2.66 | 2.63 | 1.60 | 0.58 | 2.92 | 2.76 | 2.69 | 2.89 | 2.68 | 1.38 | 0.25 | 2.56 | 2.04 |
| Ceftazoxime (ZOX) | 0.99 | 1.11 | 1.70 | 2.07 | 0.81 | 1.12 | 1.89 | 1.17 | 2.14 | 2.01 | 2.68 | 1.10 | 1.11 | 0.68 | 2.69 | 2.59 | 1.62 |
| Ceftriaxone (CRO) | 1.09 | 0.83 | 2.88 | 2.55 | 0.29 | 1.41 | 3.17 | 0.54 | 2.73 | 0.66 | 3.04 | 2.74 | 0.75 | 1.15 | 0.44 | 3.23 | 1.72 |
| Cefuroxime (CXM) | 1.75 | 0.42 | 2.94 | 2.07 | 1.70 | 2.02 | 1.91 | 1.58 | 2.92 | 2.17 | 1.94 | 1.59 | 1.68 | 2.75 | 3.27 | 2.92 | 2.10 |
| Piperacillin + tazobactam (TZP) | 2.68 | 2.71 | 3.04 | 2.43 | 2.91 | 2.45 | 0.17 | 2.50 | 2.53 | 3.31 | 0.14 | 0.61 | 2.74 | 0.09 | 0.14 | 0.17 | 1.79 |
| mean | 1.67 | 1.33 | 2.25 | 2.04 | 1.65 | 1.59 | 1.82 | 1.39 | 2.12 | 1.99 | 1.97 | 1.74 | 1.13 | 1.73 | 1.18 | 1.13 |  |

**Figure S2:**
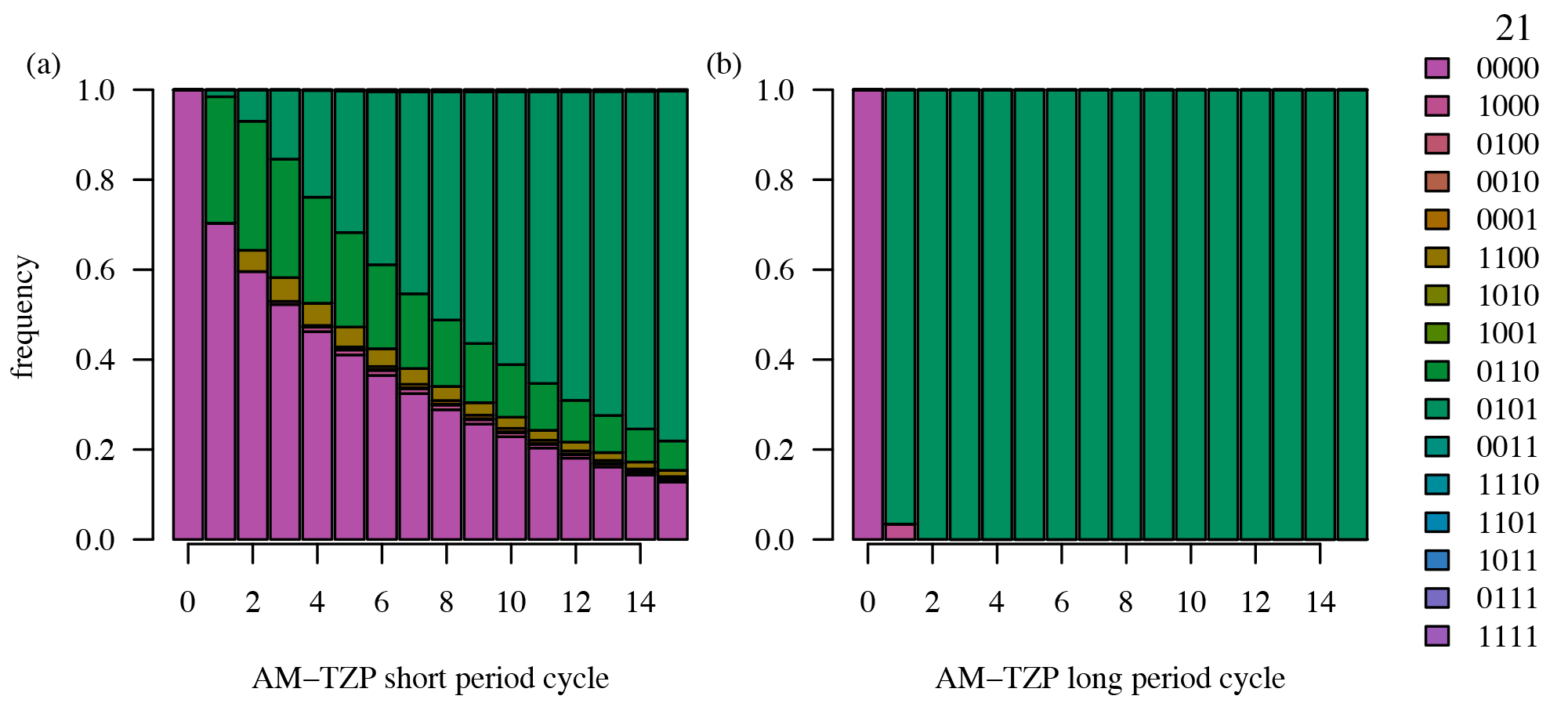
Evolution after multiple AM-TZP cycles with short (left) and long (right) periods. AM-TZP is the optimal cycle identified with a single cycle and short period [9].

**Table S2:** Optimal sequences selected by maximizing the frequency of the sensitive genotype (0000) after one cycle for multiple sequence lengths *k* under different scenarios without immigration. The growth rate refers to the long-term average growth rate after repeated cycles.

| $k$ | antibiotic sequence | frequency | growth rate ($\times 10^{-3}$) |
| --- | --- | --- | --- |
| **single cycle, short period** |  |  |  |
| 2 | AM-TZP | 0.70 | 2.72 |
| 3 | AM-TZP-FEP | 0.70 | 2.81 |
| 4 | CEC-AMC-AM-FEP | 0.62 | 2.34 |
| 5 | CEC-AMC-CPD-TZP-AM | 0.62 | 2.32 |
| **infinite cycle, short period** |  |  |  |
| 2 | CXM-AM | 0.08 | 2.37 |
| 3 | CPR-CRO-AM | 0.47 | 2.20 |
| 4 | AMC-TZP-CRO-AM | 0.58 | 2.38 |
| 5 | CTX-CXM-TZP-CRO-AM | 0.60 | 2.50 |
| **single cycle, long period** |  |  |  |
| 2 | CTT-FEP | 0.15 | 3.11 |
| 3 | CEC-AMC-FEP | 0.15 | 2.44 |
| 4 | CEC-AMC-CRO-FEP | 0.15 | 2.61 |
| 5 | CEC-AMC-CRO-CTT-FEP | 0.15 | 2.68 |
| **infinite cycle, long period** |  |  |  |
| 2 | CTT-FEP | 0.02 | 3.11 |
| 3 | TZP-CRO-FEP | 0.13 | 2.83 |
| 4 | AMC-TZP-CRO-FEP | 0.14 | 2.60 |
| 5 | CPR-CRO-AMC-CEC-FEP | 0.14 | 2.32 |

**Figure S3:**
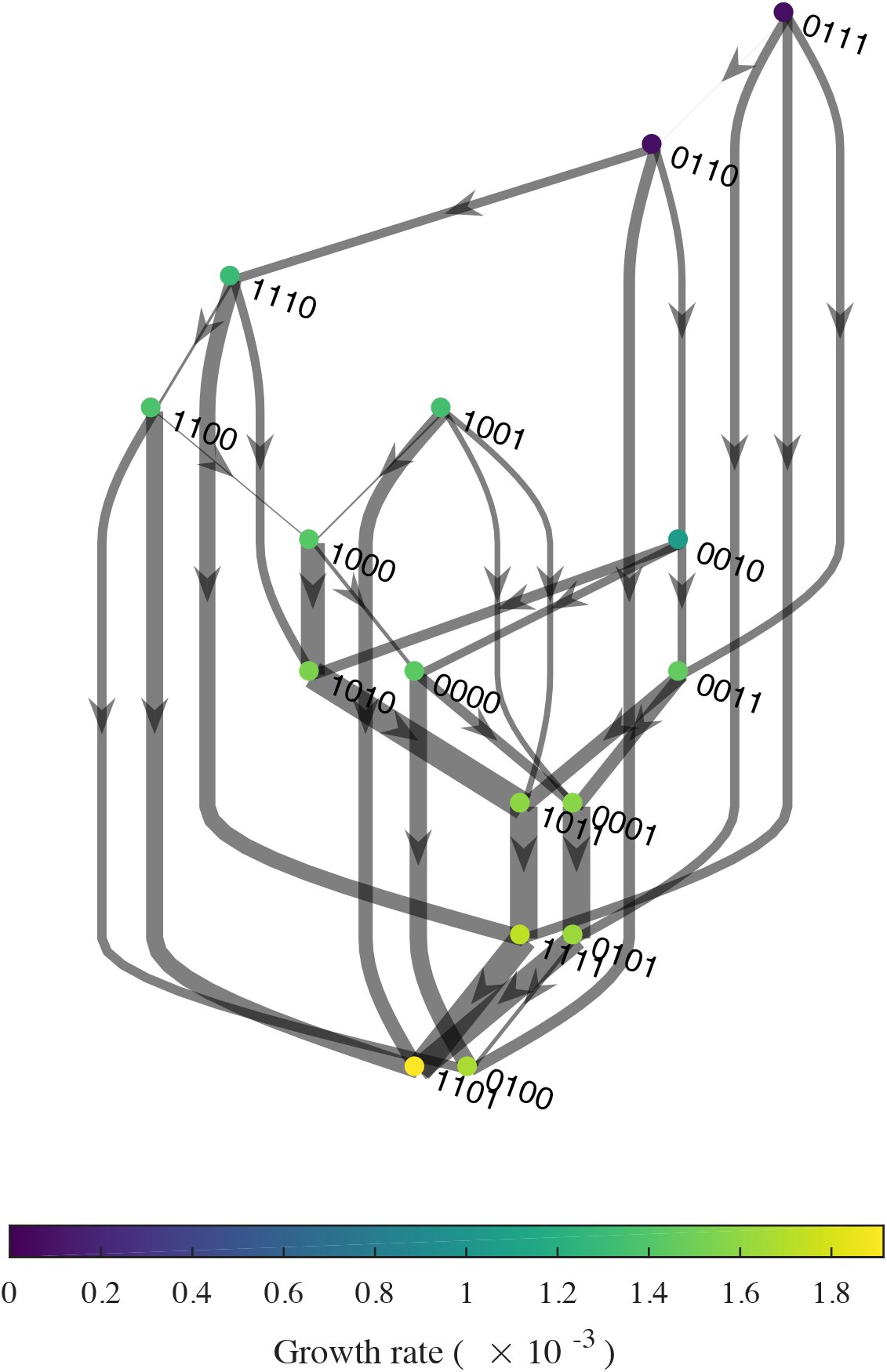
Fitness graph AMC. This figure shows the same information as Fig. 1, but the absorbing states 1101 and 0100 are more clearly depicted. Node colors show genotype growth rates. Edge weights show the relative transition probabilities *p_u,v_* from each source node *u*. Edges are not shown for transitions with zero probability. Self-loops (*p_u,u_* = 1) are omitted for clarity.

**Figure S4:**
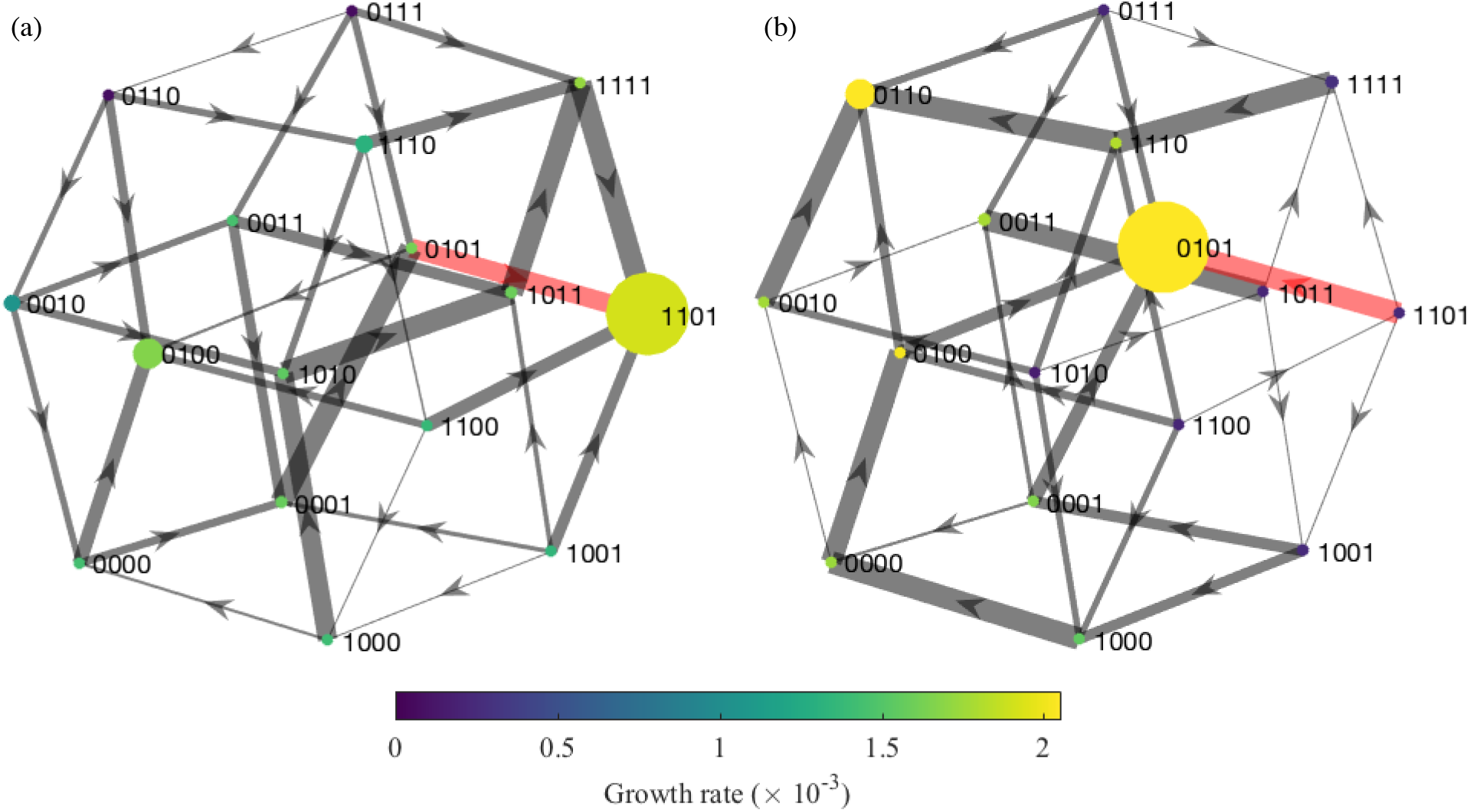
Fitness graphs during alternating (a) AMC and (b) CPR treatments for the two-drug cycle at equilibrium. Node sizes show the relative frequencies of each genotype under each drug. (Genotypes with near-zero frequencies are given visible mass to reveal color.) As in Fig. 1, node colors show genotype growth rates in each drug, and edge weights show the relative transition probabilities from each source node. The major transition between the 1101 and 0101 genotypes is highlighted in red.

**Table S3:** Optimal sequences selected by maximizing the frequency of the sensitive genotype (0000) for different values of *k* under different scenarios with immigration. The growth rate refers to the long-term average growth rate over the cycle, which was calculated by evolving the population for 300 steps under each cycle and averaging the growth rates of the last 100 steps.

| $k$ | antibiotic sequence | frequency | growth rate ($\times 10^{-3}$) |
| --- | --- | --- | --- |
| **single cycle, short period** |  |  |  |
| 2 | AM-TZP | 0.70 | 2.70 |
| 3 | AM-TZP-FEP | 0.69 | 2.78 |
| 4 | CEC-CXM-CRO-AM | 0.60 | 2.48 |
| 5 | CEC-CPD-TZP-CRO-AM | 0.60 | 2.55 |
| **infinite cycle, short period** |  |  |  |
| 2 | CXM-AM | 0.09 | 2.37 |
| 3 | CPR-CRO-AM | 0.46 | 2.20 |
| 4 | AMC-TZP-CRO-AM | 0.55 | 2.37 |
| 5 | CTX-CXM-TZP-CRO-AM | 0.57 | 2.49 |
| **single cycle, long period** |  |  |  |
| 2 | CTT-FEP | 0.10 | 3.10 |
| 3 | CRO-CTT-FEP | 0.10 | 3.03 |
| 4 | CEC-CRO-CTT-FEP | 0.10 | 2.92 |
| 5 | CEC-AMC-CRO-CTT-FEP | 0.15 | 2.67 |
| **infinite cycle, long period** |  |  |  |
| 2 | CTT-FEP | 0.10 | 3.10 |
| 3 | CEC-CTT-FEP | 0.10 | 2.97 |
| 4 | CEC-CRO-CTT-FEP | 0.10 | 2.92 |
| 5 | CPR-CRO-AMC-CEC-FEP | 0.14 | 2.32 |

**Table S4:**
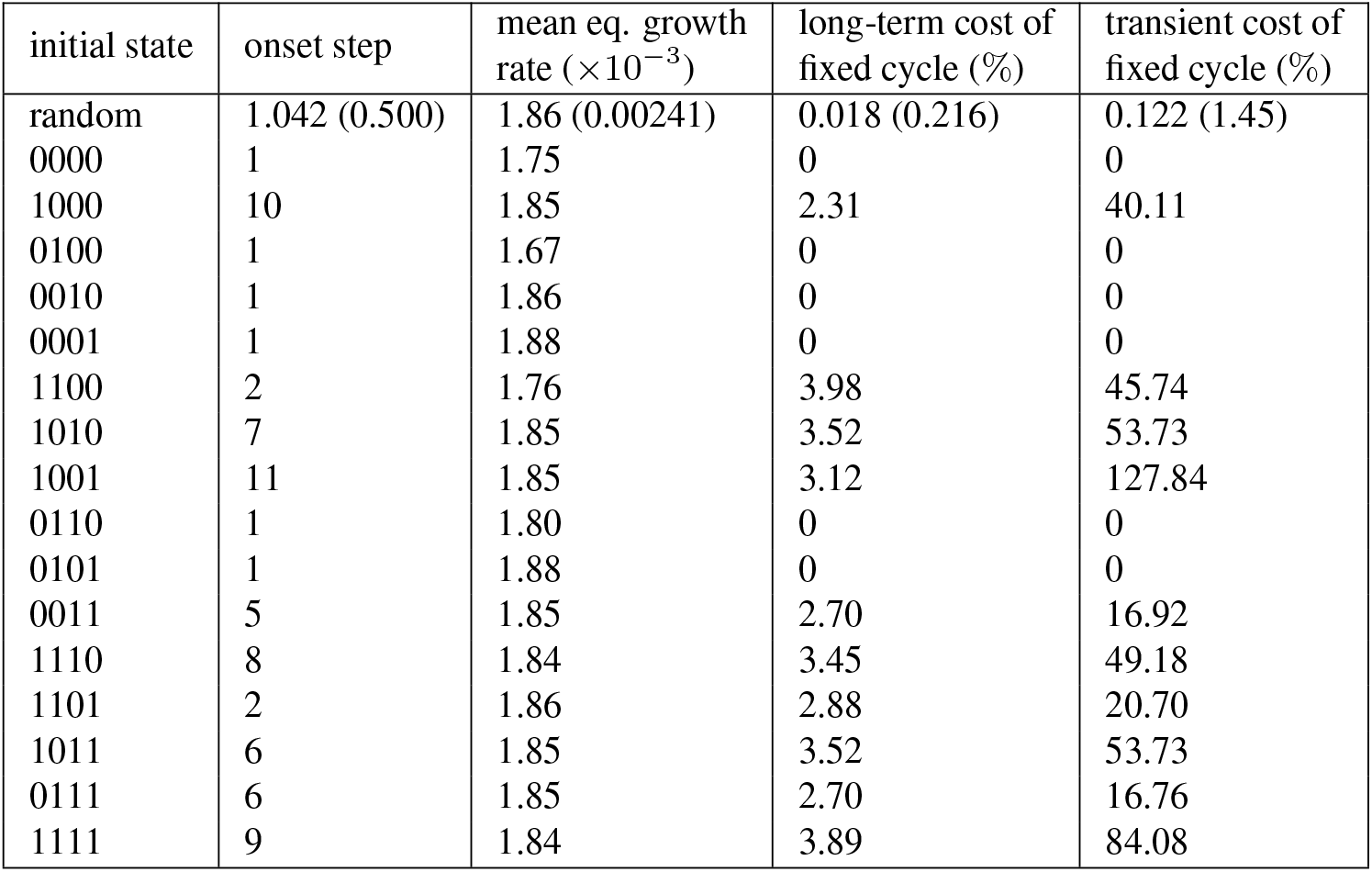
Features of AMC monotherapy, the long-term outcome of the adaptive strategy, from different starting conditions. The adaptive strategy was simulated for 300 steps starting from 1000 random initial genotype distributions and each of the genotypes individually. The onset step shows when the AMC cycles began. The mean equilibrium growth rate was calculated from the last 100 steps. The long-term cost equals the difference between the equilibrium growth rates of fixed AMC therapy and AMC therapy under the adaptive strategy. The transient cost is the cumulative sum of the differences between the growth rates under the two strategies until they converge to their stationary values. The costs (and standard deviations of costs) are reported as a percentage of the equilibrium growth rate under the corresponding initial state.

| initial state | onset step | mean eq. growth rate ($\times 10^{-3}$) | long-term cost of fixed cycle (%) | transient cost of fixed cycle (%) |
| --- | --- | --- | --- | --- |
| random | 1.042 (0.500) | 1.86 (0.00241) | 0.018 (0.216) | 0.122 (1.45) |
| 0000 | 1 | 1.75 | 0 | 0 |
| 1000 | 10 | 1.85 | 2.31 | 40.11 |
| 0100 | 1 | 1.67 | 0 | 0 |
| 0010 | 1 | 1.86 | 0 | 0 |
| 0001 | 1 | 1.88 | 0 | 0 |
| 1100 | 2 | 1.76 | 3.98 | 45.74 |
| 1010 | 7 | 1.85 | 3.52 | 53.73 |
| 1001 | 11 | 1.85 | 3.12 | 127.84 |
| 0110 | 1 | 1.80 | 0 | 0 |
| 0101 | 1 | 1.88 | 0 | 0 |
| 0011 | 5 | 1.85 | 2.70 | 16.92 |
| 1110 | 8 | 1.84 | 3.45 | 49.18 |
| 1101 | 2 | 1.86 | 2.88 | 20.70 |
| 1011 | 6 | 1.85 | 3.52 | 53.73 |
| 0111 | 6 | 1.85 | 2.70 | 16.76 |
| 1111 | 9 | 1.84 | 3.89 | 84.08 |

**Table S5:**
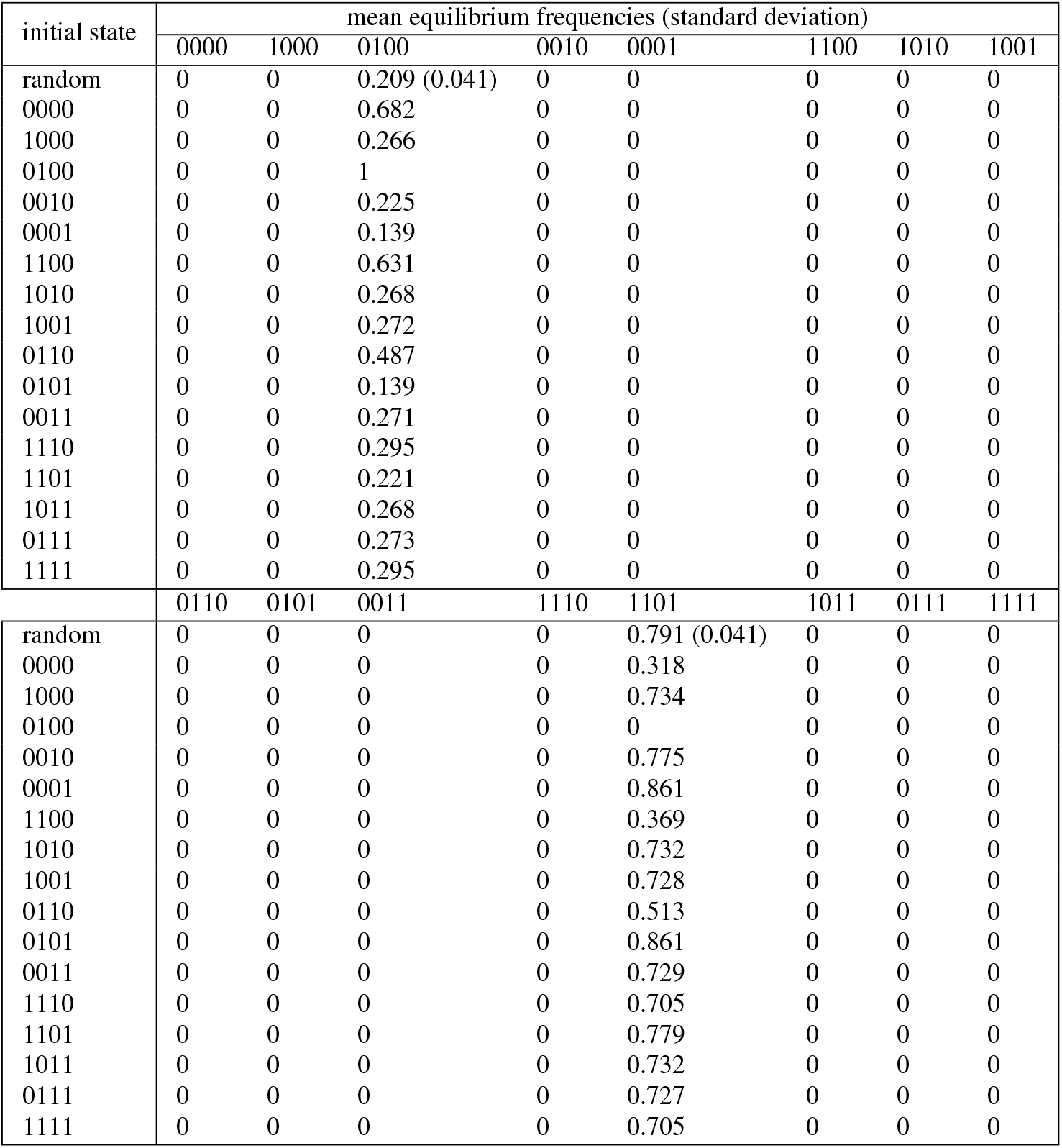
Mean equilibrium genotype frequencies with adaptive cycling starting from 1000 random initial genotype distributions and each individual genotype. The standard deviation is 0 if not given.

| initial state | mean equilibrium frequencies (standard deviation) |  |  |  |  |  |  |  |
| --- | --- | --- | --- | --- | --- | --- | --- | --- |
|  | 0000 | 1000 | 0100 | 0010 | 0001 | 1100 | 1010 | 1001 |
| random | 0 | 0 | 0.209 (0.041) | 0 | 0 | 0 | 0 | 0 |
| 0000 | 0 | 0 | 0.682 | 0 | 0 | 0 | 0 | 0 |
| 1000 | 0 | 0 | 0.266 | 0 | 0 | 0 | 0 | 0 |
| 0100 | 0 | 0 | 1 | 0 | 0 | 0 | 0 | 0 |
| 0010 | 0 | 0 | 0.225 | 0 | 0 | 0 | 0 | 0 |
| 0001 | 0 | 0 | 0.139 | 0 | 0 | 0 | 0 | 0 |
| 1100 | 0 | 0 | 0.631 | 0 | 0 | 0 | 0 | 0 |
| 1010 | 0 | 0 | 0.268 | 0 | 0 | 0 | 0 | 0 |
| 1001 | 0 | 0 | 0.272 | 0 | 0 | 0 | 0 | 0 |
| 0110 | 0 | 0 | 0.487 | 0 | 0 | 0 | 0 | 0 |
| 0101 | 0 | 0 | 0.139 | 0 | 0 | 0 | 0 | 0 |
| 0011 | 0 | 0 | 0.271 | 0 | 0 | 0 | 0 | 0 |
| 1110 | 0 | 0 | 0.295 | 0 | 0 | 0 | 0 | 0 |
| 1101 | 0 | 0 | 0.221 | 0 | 0 | 0 | 0 | 0 |
| 1011 | 0 | 0 | 0.268 | 0 | 0 | 0 | 0 | 0 |
| 0111 | 0 | 0 | 0.273 | 0 | 0 | 0 | 0 | 0 |
| 1111 | 0 | 0 | 0.295 | 0 | 0 | 0 | 0 | 0 |
|  | 0110 | 0101 | 0011 | 1110 | 1101 | 1011 | 0111 | 1111 |
| random | 0 | 0 | 0 | 0 | 0.791 (0.041) | 0 | 0 | 0 |
| 0000 | 0 | 0 | 0 | 0 | 0.318 | 0 | 0 | 0 |
| 1000 | 0 | 0 | 0 | 0 | 0.734 | 0 | 0 | 0 |
| 0100 | 0 | 0 | 0 | 0 | 0 | 0 | 0 | 0 |
| 0010 | 0 | 0 | 0 | 0 | 0.775 | 0 | 0 | 0 |
| 0001 | 0 | 0 | 0 | 0 | 0.861 | 0 | 0 | 0 |
| 1100 | 0 | 0 | 0 | 0 | 0.369 | 0 | 0 | 0 |
| 1010 | 0 | 0 | 0 | 0 | 0.732 | 0 | 0 | 0 |
| 1001 | 0 | 0 | 0 | 0 | 0.728 | 0 | 0 | 0 |
| 0110 | 0 | 0 | 0 | 0 | 0.513 | 0 | 0 | 0 |
| 0101 | 0 | 0 | 0 | 0 | 0.861 | 0 | 0 | 0 |
| 0011 | 0 | 0 | 0 | 0 | 0.729 | 0 | 0 | 0 |
| 1110 | 0 | 0 | 0 | 0 | 0.705 | 0 | 0 | 0 |
| 1101 | 0 | 0 | 0 | 0 | 0.779 | 0 | 0 | 0 |
| 1011 | 0 | 0 | 0 | 0 | 0.732 | 0 | 0 | 0 |
| 0111 | 0 | 0 | 0 | 0 | 0.727 | 0 | 0 | 0 |
| 1111 | 0 | 0 | 0 | 0 | 0.705 | 0 | 0 | 0 |

**Table S6:** Mean equilibrium genotype frequencies with fixed AMC monotherapy starting from 1000 random initial genotype distributions and each individual genotype. The standard deviation is 0 if not given.

| initial state | mean equilibrium frequencies (standard deviation) |  |  |  |  |  |  |  |
| --- | --- | --- | --- | --- | --- | --- | --- | --- |
|  | 0000 | 1000 | 0100 | 0010 | 0001 | 1100 | 1010 | 1001 |
| random | 0 | 0 | 0.207 (0.041) | 0 | 0 | 0 | 0 | 0 |
| 0000 | 0 | 0 | 0.682 | 0 | 0 | 0 | 0 | 0 |
| 1000 | 0 | 0 | 0.090 | 0 | 0 | 0 | 0 | 0 |
| 0100 | 0 | 0 | 1 | 0 | 0 | 0 | 0 | 0 |
| 0010 | 0 | 0 | 0.225 | 0 | 0 | 0 | 0 | 0 |
| 0001 | 0 | 0 | 0.139 | 0 | 0 | 0 | 0 | 0 |
| 1100 | 0 | 0 | 0.342 | 0 | 0 | 0 | 0 | 0 |
| 1010 | 0 | 0 | 0 | 0 | 0 | 0 | 0 | 0 |
| 1001 | 0 | 0 | 0.034 | 0 | 0 | 0 | 0 | 0 |
| 0110 | 0 | 0 | 0.487 | 0 | 0 | 0 | 0 | 0 |
| 0101 | 0 | 0 | 0.139 | 0 | 0 | 0 | 0 | 0 |
| 0011 | 0 | 0 | 0.065 | 0 | 0 | 0 | 0 | 0 |
| 1110 | 0 | 0 | 0.033 | 0 | 0 | 0 | 0 | 0 |
| 1101 | 0 | 0 | 0 | 0 | 0 | 0 | 0 | 0 |
| 1011 | 0 | 0 | 0 | 0 | 0 | 0 | 0 | 0 |
| 0111 | 0 | 0 | 0.067 | 0 | 0 | 0 | 0 | 0 |
| 1111 | 0 | 0 | 0 | 0 | 0 | 0 | 0 | 0 |
|  | 0110 | 0101 | 0011 | 1110 | 1101 | 1011 | 0111 | 1111 |
| random | 0 | 0 | 0 | 0 | 0.793 (0.041) | 0 | 0 | 0 |
| 0000 | 0 | 0 | 0 | 0 | 0.318 | 0 | 0 | 0 |
| 1000 | 0 | 0 | 0 | 0 | 0.910 | 0 | 0 | 0 |
| 0100 | 0 | 0 | 0 | 0 | 0 | 0 | 0 | 0 |
| 0010 | 0 | 0 | 0 | 0 | 0.775 | 0 | 0 | 0 |
| 0001 | 0 | 0 | 0 | 0 | 0.861 | 0 | 0 | 0 |
| 1100 | 0 | 0 | 0 | 0 | 0.658 | 0 | 0 | 0 |
| 1010 | 0 | 0 | 0 | 0 | 1 | 0 | 0 | 0 |
| 1001 | 0 | 0 | 0 | 0 | 0.966 | 0 | 0 | 0 |
| 0110 | 0 | 0 | 0 | 0 | 0.513 | 0 | 0 | 0 |
| 0101 | 0 | 0 | 0 | 0 | 0.861 | 0 | 0 | 0 |
| 0011 | 0 | 0 | 0 | 0 | 0.935 | 0 | 0 | 0 |
| 1110 | 0 | 0 | 0 | 0 | 0.967 | 0 | 0 | 0 |
| 1101 | 0 | 0 | 0 | 0 | 1 | 0 | 0 | 0 |
| 1011 | 0 | 0 | 0 | 0 | 1 | 0 | 0 | 0 |
| 0111 | 0 | 0 | 0 | 0 | 0.933 | 0 | 0 | 0 |
| 1111 | 0 | 0 | 0 | 0 | 1 | 0 | 0 | 0 |

